# CACTUS: integrating clonal architecture with genomic clustering and transcriptome profiling of single tumor cells

**DOI:** 10.1101/2020.06.05.134452

**Authors:** Shadi Darvish Shafighi, Szymon M Kiełbasa, Julieta Sepúlveda-Yáñez, Ramin Monajemi, Davy Cats, Hailiang Mei, Roberta Menafra, Susan Kloet, Hendrik Veelken, Cornelis A.M. van Bergen, Ewa Szczurek

**Affiliations:** Faculty of Mathematics, Informatics, and Mechanics, University of Warsaw, Stefana Banacha 2, 02-097, Warsaw, Poland; Department of Biomedical Data Sciences, Leiden University Medical Center, Einthovenweg 20, 2333 ZC, Leiden, The Netherlands; Department of Hematology, Leiden University Medical Center, Albinusdreef 2, 2333 ZA, Leiden, The Netherlands; Leiden Genome Technology Center, Leiden University Medical Center, Einthovenweg 20, 2333 ZC, Leiden, The Netherlands

**Keywords:** Single cell sequencing, Follicular lymphoma, B cell receptor, Clonal evolution, Somatic mutations, Probabilistic graphical model

## Abstract

**Background:** Drawing genotype-to-phenotype maps in tumors is of paramount importance for understanding tumor heterogeneity. Assignment of single cells to their tumor clones of origin can be approached by matching the genotypes of the clones to the mutations found in RNA sequencing of the cells. The confidence of the cell-to-clone mapping can be increased by accounting for additional measurements. Follicular lymphoma, a malignancy of mature B cells that continuously acquire mutations in parallel in the exome and in B-cell receptor loci, presents a unique opportunity to align exome-derived mutations with B-cell receptor clonotypes as an independent measure for clonal evolution.

**Results:** Here, we propose CACTUS, a probabilistic model that leverages the information from an independent genomic clustering of cells and exploits the scarce single cell RNA sequencing data to map single cells to given imperfect genotypes of tumor clones. We apply CACTUS to two follicular lymphoma patient samples, integrating three measurements: whole exome sequencing, single cell RNA sequencing, and B-cell receptor sequencing. CACTUS outperforms a predecessor model by confidently assigning cells and B-cell receptor clonotypes to the tumor clones.

**Conclusions:** The integration of independent measurements increases model certainty and is the key to improving model performance in the challenging task of charting the genotype-to-phenotype maps in tumors. CACTUS opens the avenue to study the functional implications of tumor heterogeneity, and origins of resistance to targeted therapies.

## Introduction

Tumor heterogeneity and clonal evolution present a major challenge for cancer therapy (1). Tumor cells carry founder and subsequently acquired driver mutations that cause transformation of the healthy cell into an expanding population of malignant cells. Continuous acquisition of mutations creates populations of tumor cells with divergent mutational profiles. Diverging cells with acquired driver mutations result in preferential clonal expansion leading to intraclonal diversity. Given that distinct genotypes induce key phenotypic differences between the clones (2), considerable gene expression variation is expected between the clones. Measuring the phenotypes of tumor clones, however, is challenged by the difficulties in resolving the clonal genotype-to-phenotype maps in tumors (3). Follicular lymphoma (FL) is a common type of malignant B-cell lymphoma with characteristics of normal germinal center (GC) B-cells. FL pathogenesis is founded by the paradigmatic translocation (14;18)(q32;q21) that places BCL-2 under transcriptional control of the IGH@ locus enhancer. Secondary drivers affect genetic modifiers that enhance germinal center (GC) formation and reduce B-cell differentiation beyond the GC stage (4, 5). Despite commonly observed pathogenic genomic events, clinical behaviour of FL is unpredictable and ranges from spontaneous remission over long-term stable disease to transformation to aggressive B-cell lymphoma.

In addition, FL cells are continuously exposed to a physiological mutator mechanism, i.e. constitutive expression and action of activation induced cytidine deamidase (AID) (6). AID focuses on B-cell receptor (BCR) loci and results in highly mutated BCR heavy and light chain encoding genes (7). Whereas BCR mutations intrinsically may lead to a proliferative signal by acquisition of N-linked glycosylation (8), preferential expansion of clones with identical BCR can also be explained by underlying driver mutations that enhance proliferation to the BCR clone or group of BCR clones. In addition to grouping of individual cells into evolutionary clones by exome-wide mutations and structural variants, single FL cells can also be clustered based on the expression of identical BCR sequences, or BCR clonotypes. BCR mutations can therefore be considered events in clonal evolution in FL and present suitable markers that may allow a more accurate reconstruction of clonal evolution than based on mutations only.

Elucidation of tumor evolution and reconstruction of the tumor clonal architecture is possible from bulk DNA sequencing (9–12) and from single cell (sc) DNA sequencing data (13–16). Recent efforts into the direction of mapping genotypes to phenotypes in tumors include characterizing gene expression profiles of tumor clones based on matching the scRNA-seq readouts to copy number variants in the clones (17–19). Poirion et al. (20) proposed a linear model detecting statistical association of single nucleotide variants from scRNA-seq with gene expression. This approach, however, ignores the evolutionary history of the tumor, which can be resolved to determine the genotypes of the tumor clones, and the fact that mutations observable in scRNA-seq can be matched with the clone genotypes. Recently introduced cardelino (21) is the first approach to successfully utilize the mutation mapping between the clone genotypes and the variants in scRNA-seq data. The performance of this approach, however, can be hampered by the fact that sc transcripts contain only information on 5’ part of the RNA and that the data are sparse. To increase the confidence of clonal genotype to gene expression phenotype mapping, additional available evidence, such as the grouping of cells into BCR clonotypes in FL evolution, should be integrated into the inference. Combining multiple data sources has the potential to increase the resolution of tumor heterogeneity analysis (22), but presents a computational challenge (23) and calls for a dedicated probabilistic model.

Here, we propose a probabilistic graphical model for integrating Clonal Architecture with genomic Clustering and Transcrip-tome profiling of single tUmor cellS (CACTUS). The model extends cardelino (21) and maps single cells to their clones based on comparing the allele specific transcript counts on mutated positions to given clonal genotypes, leveraging additional information about evolutionary cell clusters. CACTUS is applied to newly generated data to assign single cells derived from two FL tumor samples to their clones of origin, accounting for their BCR clonotypes (Fig. 1). We demonstrate that guided by the BCR sequence information, CACTUS assigns single cells to tumor clones in agreement with independent gene expression clustering. For both subjects, CACTUS maps cells and BCR clonotypes with substantially higher confidence than cardelino. These results indicate that the important challenge of tumor genotype-to-phenotype mapping can successfully be approached by probabilistic integration of multiple measurements.

**Fig. 1.**
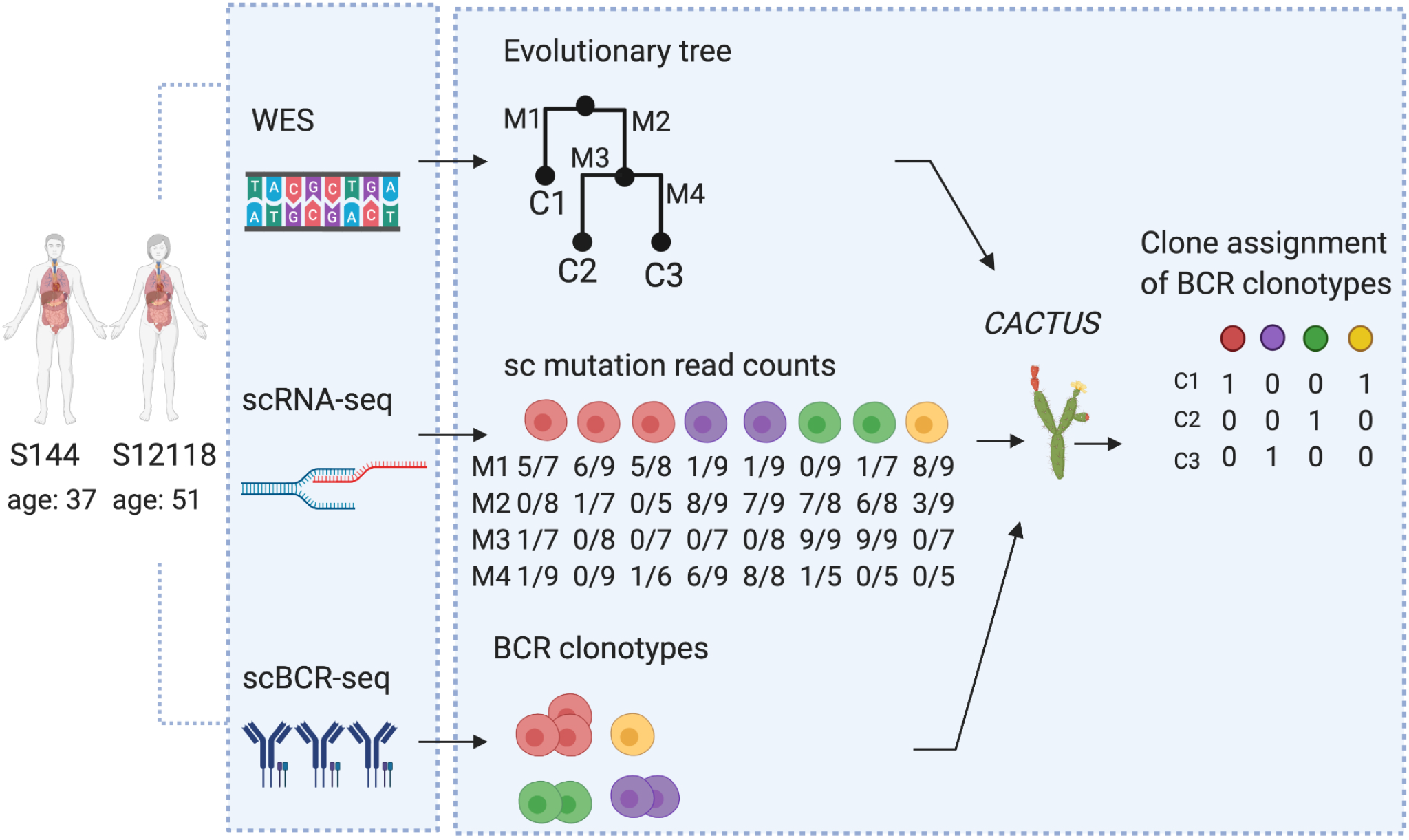
Overview of the patient data analysis and the CACTUS model. Whole exome sequencing and single-cell sequencing of all transcripts, as well as single-cell sequencing of BCR was performed on samples from two FL patients. Using WES, imperfect clonal evolution could be inferred and given as a prior to the model (C1, C2, *…*). From scRNA-seq, allele specific transcript counts (mutated/total) were extracted at mutated positions (M1,M2, *…*). BCR clonotypes were defined as clusters of cells with identical BCR heavy chain sequences. The data of input tumor clones, mutation transcript counts, and given single cell clusters (here, the BCR clonotypes) are combined in the CACTUS model for inference of the clonal assignment of the clusters.

## Results

### Single cell and WES profiling of two FL patients

The analyzed cell populations were collected from lymph nodes of two FL patients: a male patient (S144) at the age of 37, who was diagnosed with an IgM expressing FL stage IV and a female patient (S12118) at the age of 51, who was diagnosed with an IgG expressing FL stage IV. To detect (sub-)clonal mutations, we performed whole exome sequencing (WES) at 200x coverage and called mutations between FL cells and paired stromal non-hematopoietic cells. We detected 398 somatic mutations for patient S144 and 1034 somatic mutations for patient S12118 with somatic p-value (SPV) < 0.1.

Next, we performed pooled single cell sequencing of purified FL cells simultaneously for full transcriptomes and BCR enriched libraries. We used the Vireo method (24) to assign single cells to patients based on matching of alleles expressed in the single cells with mutations detected by bulk WES. Deconvolution of the whole transcriptome data yielded counts for 1524 cells of subject S144 and 874 cells of subject S12118. BCR sequencing yielded BCR heavy chain clonotypes (cells with identical BCR sequences) for approx. 70% of cells in both patients. Both samples were dominated by a limited number of large BCR clonotypes with many rare BCR clonotypes. ‘Pielou evenness index’ was 0.59 for S144 and 0.53 for S12118, indicating moderate intraclonal diversification (25). For generality, cells without BCR heavy chain sequences were considered to form a separate clonotype with only one cell (see Figure S1 for BCR clonotype size distribution).

### A probabilistic model for assigning cell clusters to evolutionary tumor clones

CACTUS is a Bayesian method that integrates three different sources of prior knowledge: a set of tumor clones with their genotypes, independently obtained non-overlapping cell clusters, and scRNA-seq transcripts at mutated sites, to map each cell cluster to its corresponding clone. Cells of the same cluster are assumed to come from the same tumor clone. Since the clusters are non-overlapping sets of cells, the cluster assignment to clones naturally defines also cell assignment (each cell in a given cluster is assigned to the same clone as its cluster). Here, the cell clustering was defined by the BCR clonotypes. Cells of the same BCR clonotype are expected to come from the same tumor clone. Thus, here CACTUS took advantage of the extra information of BCR sequences to gain power and precision of the assignment.

CACTUS yields the posterior probability estimate for each given cell cluster to be mapped to each given clone. This probability is computed using a beta-binomial model for the allele specific transcript counts for each mutation and cell in this cluster. The model estimates the error rate for the given imperfect genotypes of the clones and infers corrected genotypes. The likelihood of assigning a cluster to a given clone increases with the similarity of the mutation signal observed in the cells of the cluster to the corrected genotype of that clone. The final assignment of the clusters (and thus also their contained single cells) is defined by selecting the most probable tumor clone for each cluster (here, BCR clonotype; Fig. 1).

For both subjects, to define the input clonal structures, we first identified a set of such mutations that could be found both in WES and scRNA-seq data. From the identified 398 mutations with SPV < 0.1 for subject S144 and 1034 mutations for subject S12118, for further analysis we selected only these mutations, for which any transcript expression was observed in scRNA-seq. Despite the relaxed significance level of 0.1 for the somatic p-values, we consider the common mutations as reliable, since they have evidence in both data sources. Only 5 out of 398 total resulting common mutations for subject S144, and 5 out of 137 common mutations for subject S12118, had somatic p-value in the (0.01, 0.05) interval (Figure S2). Numbers of the common mutations vary in different cells (Figure S3). For further analysis we considered only cells which contain at least one of the common mutations. This included 1262 out of 1524 cells in subject S144 and 799 out of 874 cells in subject S12118.

We next applied Canopy to the WES data for the common mutations, and extracted the top tree and its corresponding clones, with their genotypes. To obtain the cell-to-clone assignment, CACTUS was applied to the obtained clonal structure, with BCR clonotypes and scRNA-seq transcript counts as input. To demonstrate how the addition of the BCR clonotype information improves the assignment of cells to clones, we applied cardelino (21) to the same Canopy trees and the scRNA-seq transcript counts. From these data, cardelino derived cell assignment to tumor clones. The two models (CACTUS and cardelino) are similar, but CACTUS is more general as it takes into account the cell clustering (here, BCR clonotype) information. In fact, for the specific case of such uninformative clustering that contains exactly one cell in each cluster, CACTUS reduces to cardelino. Thus, naturally, the advantage of CACTUS should be most visible for such cells that are contained in clusters of more than one cell.

### CACTUS solution verified by an independent gene expression analysis

To validate the BCR clonotype assignment and the induced cell assignment, we performed independent analysis of transcript expression levels obtained from scRNA-seq of the same cells. First, we investigated whether the grouping of cells into the inferred clones coincides with similarity of their expression profiles (Fig. 2, 3). To this end, we reduced the dimension of expression data using UMAP (26) and t-SNE (27) provided in the Seurat package (28) and colored each cell with its corresponding clone inferred using CACTUS, and for a comparison, cardelino (21).

**Fig. 2.**
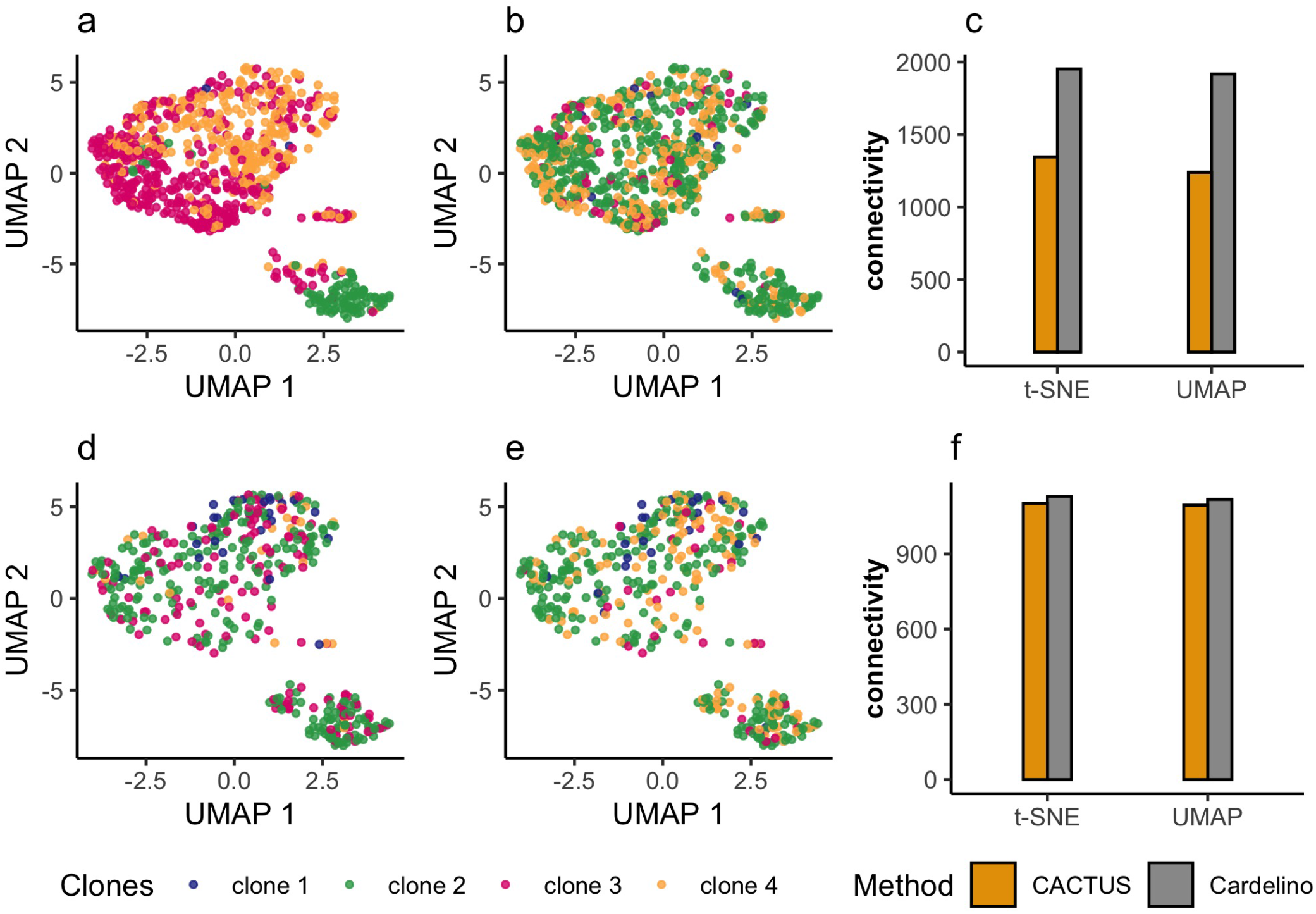
Validation of cell-to-clone assignment with gene expression for subject S144. **a, b, d, e** Transcript expression of the cells reduced to two dimensions using UMAP, shown separately for the cells in BCR clonotypes containing more than one cell (**a**, **b**) and for cells belonging to BCR clonotypes with only one cell (**d,e**). Each point corresponding to a cell is colored by its clone assigned by CACTUS (**a**, **d**) and by cardelino (21) (**b**, **e**). The low connectivity values indicate that using CACTUS, the cells in the same clone are also close in the reduced gene expression space (**c**, **f**). The advantage of CACTUS is more pronounced for cells in BCR clonotypes containing more than one cell.

**Fig. 3.**
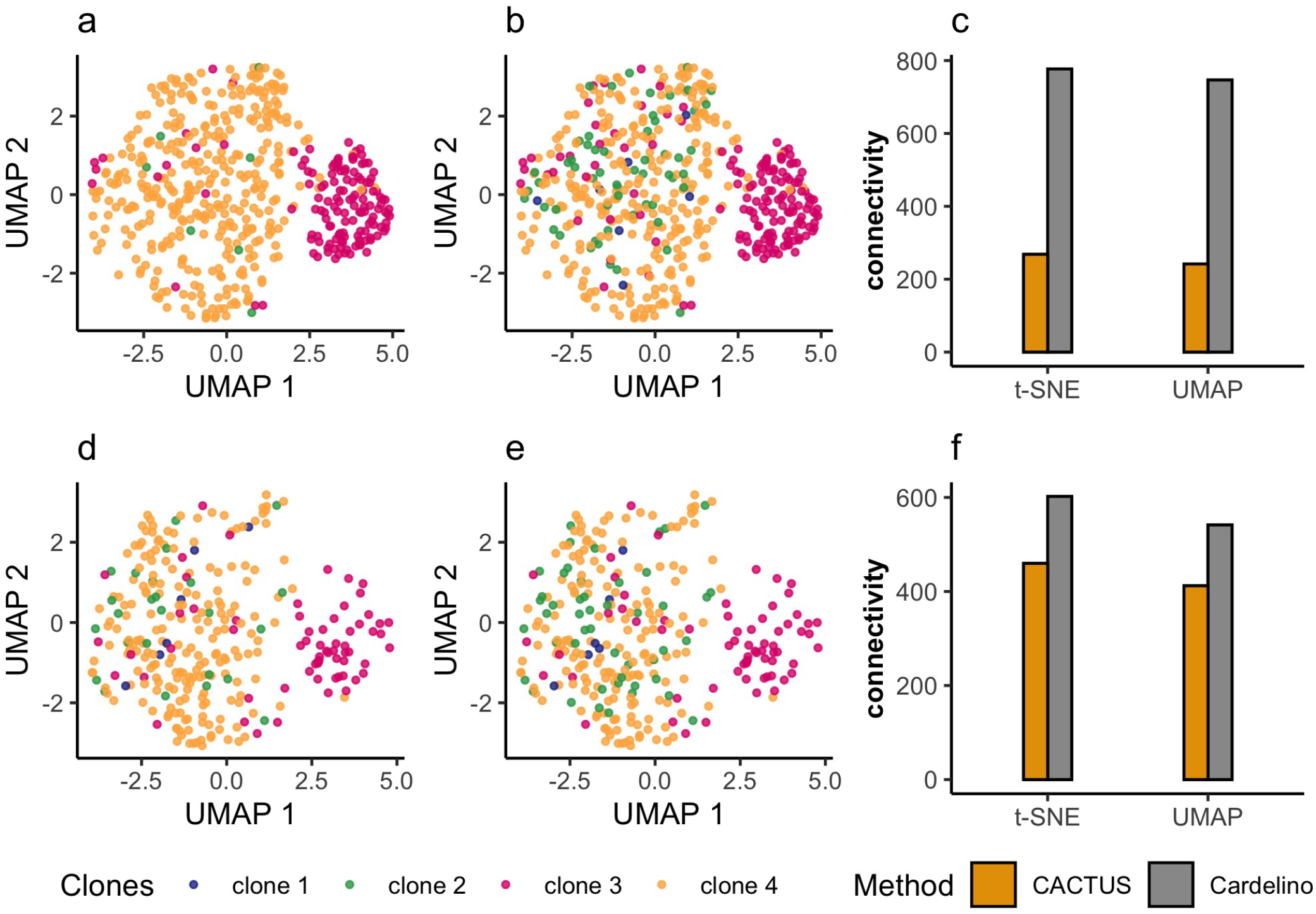
Validation of cell-to-clone assignment with gene expression for subject S12118. Figure panels as for subject S144 in Fig. 2. Also for subject S12118, assignment to clones for cells in BCR clonotypes of more than one cell using CACTUS (**a**) improves agreement with gene expression data compared to assignment of cells in BCR clonotypes containing only one cell (**d**) and assignment using cardelino (21) (**b**), as quantified using connectivity measure (**c**). For BCR clonotypes containing only one cell CACTUS performs comparably well as cardelino.

As expected, CACTUS leverages information obtained from the BCR clonotypes containing more than only one cell. For cells in such BCR clonotypes, the results of CACTUS are more consistent with gene expression (visualized for UMAP in Fig. 2a and Fig. 3a) than the results of cardelino (Fig. 2b and Fig. 3b). For subject S144 and cells contained in BCR clonotypes with more than one cell, CACTUS identifies clone 2 as a set of cells that are clearly separated in gene expression space from a large cluster of cells, which is populated in one half by clone 3 and clone 4, while cardelino finds clones which are mixed in the reduced gene expression space (Fig. 2). For subject S12118, both methods associate clone 3 with one gene expression cluster and clone 4 with another, with the two gene expression clusters clearly separated in the reduced space. For CACTUS, the identified clones are slightly less intermixed with others than for cardelino (Fig. 3). To quantify the agreement of the obtained assignment of cells to the clones with gene expression, we used a connectivity measure (29). The connectivity measure would be minimized when the cells in the same clone would also be close in terms of Euclidean distance in the reduced gene expression space. For cells in BCR clonotypes with more than one cell, CACTUS obtains significantly lower connectivity values, regardless whether t-SNE or UMAP is used for dimensionality reduction (Fig. 2c and Fig. 3c).

In contrast, clone assignments of cells that are not grouped into larger BCR clonotypes, show less agreement with gene expression (Fig. 2d and Fig. 3d). This agreement for those cells is comparably low for cardelino (Fig. 2e and Fig. 3e). Still, the connectivity values for the CACTUS results are a bit better (lower) than for cardelino (Fig. 2f and Fig. 3f).

Second, we performed independent clustering of cells by their normalised gene expression using Seurat (28). Then, we compared the resulting gene expression clusters to the clones inferred by CACTUS and by cardelino using the Adjusted Rand Index (shortly, ARI; (30)). The index is a corrected-for-chance version of the Rand index, measuring similarity between two given clusterings, with values in the [−1, 1] interval. ARI is negative when the agreement is lower than expected and is maximized when the compared clusterings are identical. For subject S144 and cells that are not grouped into larger BCR clonotypes, both clones inferred by CACTUS and by cardelino show no similarity to gene expression (with ARI 0.0002 and 0.0017, respectively). Compared to cardelino (ARI 0.0055), CACTUS achieves a higher agreement with the gene expression clustering for cells contained in clonotypes with more than one cell (ARI 0.16). For subject S12118, the CACTUS clones again show a higher similarity to gene expression clusters. For cells that are not grouped into larger BCR clonotypes, CACTUS yields ARI of 0.2, while cardelino 0.16. Finally, for cells in clonotypes of more than one cell, the ARI for CACTUS is 0.3, while for cardelino it is 0.25. Overall, these results indicate, that by accounting for the BCR clonotypes, CACTUS improves the genotype-to-gene expression phenotype mapping.

### CACTUS enhances the confidence of cell-to-clone assignment

For both subjects, the top identified evolutionary trees consisted of four clones (Fig. 4a, b). The number of mutations acquired along the branches of the trees ranges from 0 to 57. The genotype of each clone is defined as the set of the mutations acquired on the path from the root of the tree to the leaf corresponding to the clone (Table S1). Notably, the clone frequencies derived by Canopy (Fig. 4a, b) have been corrected both by CACTUS (Fig. 4c, g, e, i) and cardelino (Fig. 4d, h, f, j).

**Fig. 4.**
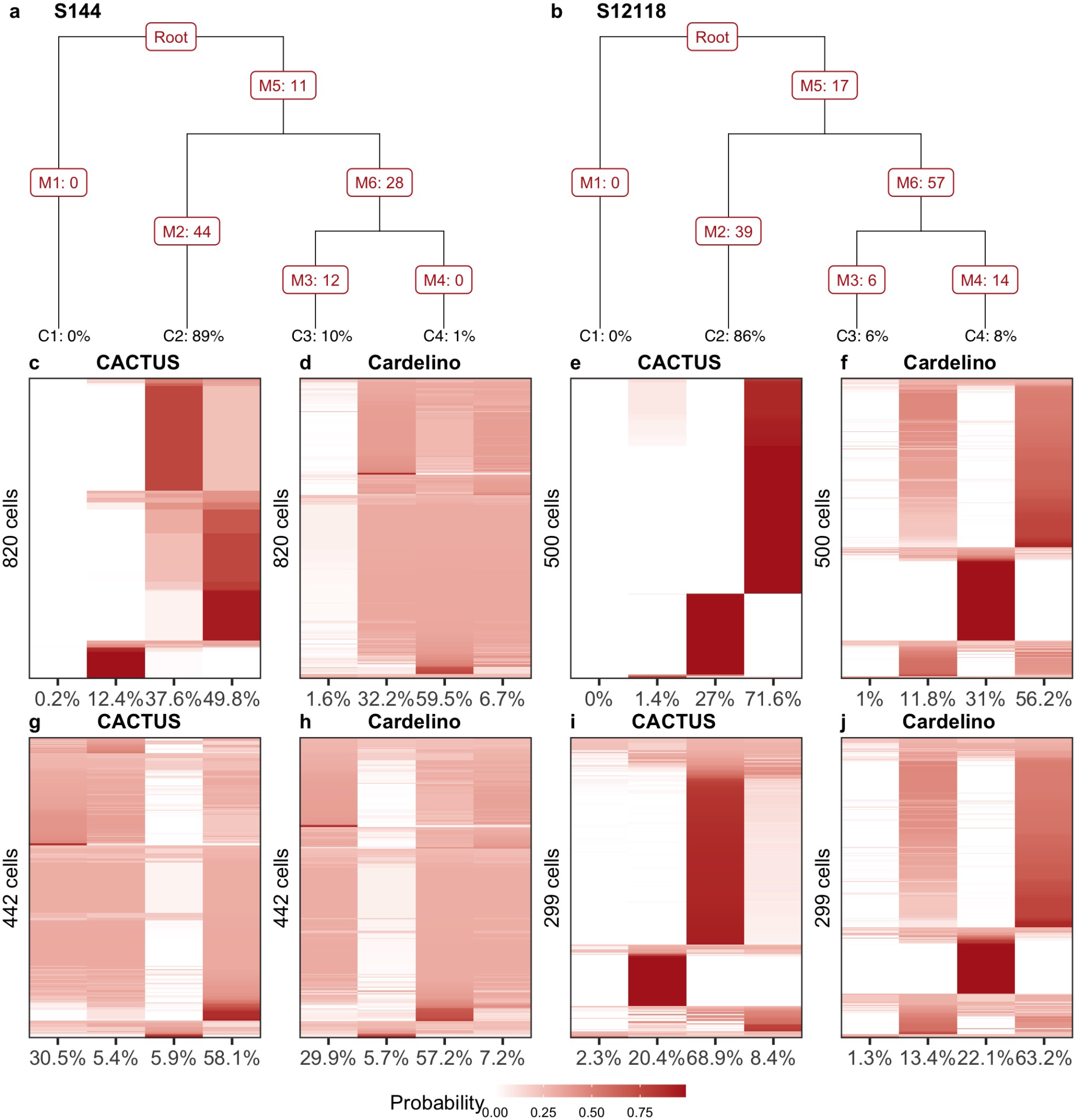
Confidence of cell assignment to the tumor clones. **a, b** Evolutionary trees inferred by Canopy (9) for subject S144 (**a**) and S12118 (**b**). Leaf labels: clone prevalences. Branch labels: numbers of acquired mutations (can be zero). Clone 1 corresponds to the base, normal clone. In tree **a**, clone 4 (C4) differs from clone 3 (C3) by the 12 SNVs acquired on the branch leading to the leaf C3. Canopy considers also CNVs, but they are not used for cell-to-clone mapping and hence not visualized here. **c-j** Shades of brown indicate the probability of assignment of cells (y axis) to the clones (x axis; labeled with corrected prevalences) by CACTUS (**c, g, e, i**) and cardelino (21) (**d, h, f, j**). For cells in BCR clonotypes with more than one cell (second row), CACTUS yields higher confidence of cell-to-clone assignment (**c, e**) than cardelino (**d, f**). For cells in BCR clonotypes with only one cell (third row) for subject S144 the confidence of cell-to-clone assignment by CACTUS (**g**) is similarly weak as by cardelino (**h**), while for S12118 and for CACTUS (**i**) the confidence is higher than for cardelino (**j**).

Next, we investigated the confidence of assignment of cells to the tumor clones for both subjects (Fig. 4). The assignment of cells to the clones was directly derived from the assignment of their BCR clonotypes. In general, thanks to the additional information from the BCR clonotypes, CACTUS assigns cells to clones with a clearly higher confidence than cardelino (21). For subject S144 and majority of cells, the probability of assignment by cardelino is almost uniform across the clones (Fig. 4d, h). In contrast, for the subset of cells in BCR clonotypes with more than one cell, CACTUS makes confident assignments (Fig. 4c). For cells in one-cell BCR clonotypes CACTUS assigns cells with similar confidence to cardelino (Fig. 4g).

Compared to S144, for subject S12118 the confidence of assignment is larger for both methods (Fig. 4). Again, CACTUS has an advantage over cardelino, especially for cells in BCR clonotypes with more than one cell, assigning a majority of those cells to one clone with high probability (Fig. 4e,i). In contrast, for majority of cells, cardelino yields similar probabilities of assignment to clones 2 and 4 (Fig. 4f, j).

### Assignment of BCR clonotypes to tumor clones

Finally, we inspected the assignment of BCR clonotypes to clones by CACTUS. For a comparison, we computed the proportion of each BCR clonotype that contained more than one cell (the fraction of cells in that BCR clonotype) that were assigned to each clone using cardelino (Fig. 5). In the case of ties in the highest proportions across clones, we assumed the BCR clonotype was assigned to the same clone as by CACTUS. As expected by construction of the underlying probabilistic model, for both subjects, CACTUS assigns entire BCR clonotypes to single clones (Fig. 5a, c). For cardelino, the proportions of BCR clonotypes are more distributed across the clones (Fig. 5b, d). Given the uncertainty of assignment of cells to clones by cardelino for subject S144 (Fig. 4), it is unsurprising that for some of the BCR clonotypes, the clone assigned by CACTUS does not agree with the clone with the highest proportion of cells assigned by cardelino. Both methods agree on the assignment of clonotype U to clone 1. All of 14 BCR clonotypes assigned to clone 2 by CACTUS, were assigned to the same clone by cardelino. Out of 17 BCR clonotypes assigned to clone 3 by CACTUS, however, only 1 is assigned to clone 3 also by cardelino. This large disagreement comes mainly from the fact that cardelino assigned the highest proportion of cells contained in 15 of these 17 clonotypes again to clone 2. Finally, out of 5 BCR clonotypes assigned to clone 4 by CACTUS, 2 were assigned in the highest proportion to the same clone also by cardelino.

**Fig. 5.**
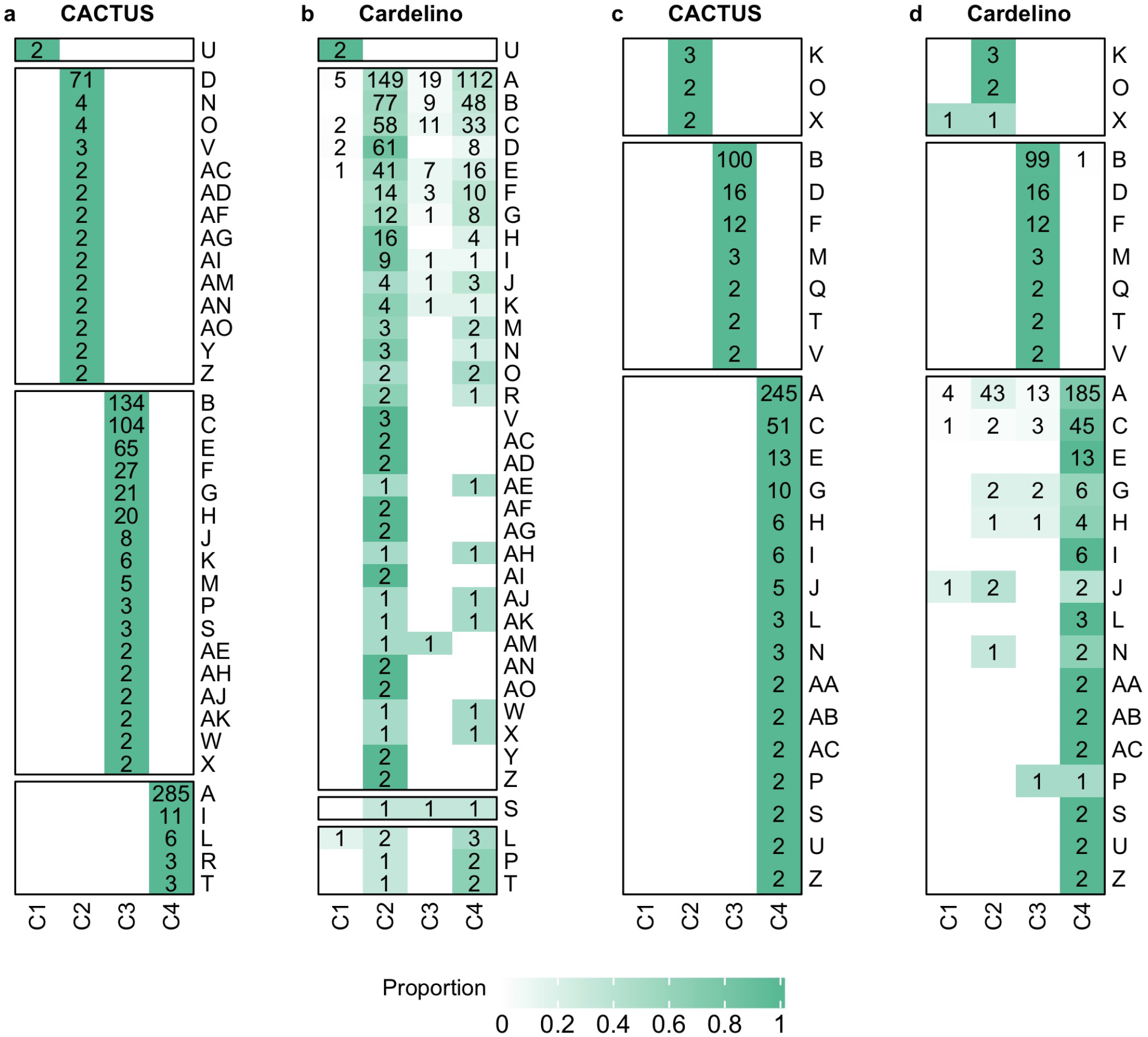
**BCR clonotype assignment to tumor clones,** for both subjects: S144 (**a, b**) and S12118 (**c, d**), using CACTUS (**a, c**) and cardelino (21) (**b, d**). Heatmaps with shades of green indicate the proportion of cells in clonotypes (y axis) assigned to clones (x axis). Each number in a green entry indicates the nonzero number of cells of the corresponding BCR clonotype assigned to the corresponding clone. Only BCR clonotypes of at least two cells are featured. As expected, for both subjects, CACTUS assigns entire BCR clonotypes to single clones (**a, c**). For cardelino, the proportions of BCR clonotypes are more distributed across the clones (**b, d**).

For subject S12118, the assignment of clonotypes agrees between the two methods. This is in accordance with the increased confidence of assignment of cells to clones by both methods for that subject (compare Fig. 4).

In summary, the agreement of both cell-to-clone and BCR clonotype-to-clone mapping between the CACTUS and cardelino increases with the confidence of assignment. For subject S144, for which cardelino yielded low-confidence assignments, 746 out of 1262 cells in total (59%) and 19 out of 37 BCR clonotypes with more than one cell (51%) were assigned to different clones by the two methods. Here, we assume cardelino assigns a BCR clonotype to the clone to which it assigned the highest proportion of cells. For subject S12118, where both methods increased confidence of assignment, only 144 cells out of 799 (14%) and no BCR clonotypes out of 26 clonotypes with more than one cell were assigned differently.

## Discussion

Here, we propose a probabilistic model for accurate and confident mapping of single tumor cells to their evolutionary clones of origin. In this way, it allows clone-specific gene expression profiling, opening the possibility to reconstruct genotype-to-phenotype maps. The task of cell-to-clone mapping is challenged by multiple technical obstacles. First, although multiple methods exist for the inference of tumor evolution, resolving tumor clones and their genotypes is in itself a difficult computational problem and errors are expected (12). Thus, CACTUS, uses the additional signal both in the scRNA-seq and in clustering data to correct the given genotypes of the clones. Second, the information in scRNA-seq data is only sparse, prone to errors such as dropout and uneven coverage, and biased to mutations observable in typically sequenced 150 bp of transcripts. It is thus important to realize that the analysed tumor history is limited only to the mutations measurable in single cells, and is potentially more coarse-grained than the true clonal structure of the tumor. These limitation are purely technical, and in this respect analysis using CACTUS would benefit from full-length transcript sequencing with high depth, as well as further developments increasing the quality of scRNA-seq technology.

The key aspect of our model is the ability to borrow statistical strength across different measurements (both of DNA and RNA) of the cells in the sample. In particular, in addition to clone genotypes derived from WES, and allele specific read counts measured using scRNA-seq, the model leverages information given by cell clustering. Our results show that this additional evidence is crucial to overcome the challenges of the cell-to-clone assignment problem. Not any given cell clustering, however, can empower CACTUS to deliver more confident results. The assumption that cells contained in the same cluster belong to the same clone is critical for model performance. In particular, such cell clustering, where the cells in the same cluster are not expected to belong to the same clone, can misguide model inference. Apart from clustering by genomic features, which is expected to agree with the clonal structure of the tumor cell population, for example, clustering by location in the tissue could be provided as input to CACTUS. Here, we used BCR clonotypes to define the clustering. As would other relevant genomic features, mutations in BCR loci bring evolutionary information. On a general level, they indicate whether a BCR clonotype is relatively old with a low number of BCR mutations, or if a BCR clonotype has more recently evolved and carries a higher number of mutations. Identical BCR sequences indicate common evolutionary origin, as otherwise they would be disrupted by acquisition of additional mutations. The quality of additional information brought in by the BCR clonotypes is assured by the complete and deep sequencing coverage of BCR loci in the applied scRNA-seq strategy.

CACTUS could be extended in the future to further broaden its functionality and to account for even more additional measurements. First, at the moment the model considers the given cell clustering as error-free, i.e., that each cell is assigned to the correct cluster. This assumption could be relaxed, and the model could utilize the signal in the other input data to correct the potential clustering errors. The input genotypes and the number of clones are corrected, but need to be inferred *a priori* to applying the model. Instead, CACTUS could be extended to simultaneously infer the evolutionary tree, yielding the clones and their genotypes, together with the cell assignment to the clones. Finally, other measurements could be incorporated to statistically strengthen model inference. For example, gene expression similarities between cells, here used for model validation, could be used as input, as cells with similar expression profiles are expected to come from the same clone.

The model is applied to newly generated FL patient data, for the first time shedding light on how clonal evolution in this cancer type induces clone-specific gene expression and agrees with BCR clonotypes. Accurate mapping of clonal structures with gene expression patterns allows detection of potential therapy-resistant clones, which is essential for effective personalized treatment. Our results demonstrate applicability of CACTUS to the complex cancer samples. The model, however, is more generally applicable and can describe somatic evolution also in other diseases or in the healthy tissue.

## Conclusions

Here, we deal with the task of gene expression profiling of tumor clones by matching the genotypes of the clones to the mutations found in RNA sequencing of the cells. CACTUS benefits from the additional information contained in the BCR clonotypes to assign cells to clones, to successfully deal with errors and dropouts in single cell RNA sequencing, and the difficulty of inferring the correct clonal structure. In summary, this contribution is a step forward in establishing computational tools for resolving the tumor heterogeneity and, by combining genotype with gene expression profiles, its impact on functional diversification of the tumor cell subpopulations.

## Methods

### Follicular Lymphoma sample preparation

Samples with histologically confirmed infiltration of follicular lymphoma were collected with approval by the institutional review board of Leiden University Medical Center according to the declaration of Helsinki and with written informed consent. Single cell suspensions were obtained by gentle mechanical disruption and mesh filtration and were cryopreserved using 10% DMSO as cryoprotectant. Remaining tissue was cultured in low-glucose DMEM to obtain stromal cell cultures for isolation of DNA of nonmalignant cells. Thawed single FL cells were purified by flow cytometry using fluorescently labeled antibodies specific for CD19 and CD10 and rested overnight followed by removal of dead cells using MACS dead cell removal kit. Cells of different patients were pooled and loaded on a 10X Genomics chip to obtain single cell cDNA libraries for an expected 1500 cells per patient. Following single cell cDNA library generation and amplification, one fraction was directly sequenced for 5’ gene expression profiling. The second fraction was enriched for BCR transcripts by seminested amplification using 3’ constant domain primers for all BCR genes, partially digested and sequenced. Both single cell libraries were sequenced in paired-end mode on Illumina (2×150 bp).

### WES sequencing and mutation calling

FL single cells were purified by flow cytometry as described above to obtain bulk purified FL cells for immediate isolation of DNA. Whole exome sequencing (WES) was performed on paired FL and normal DNA at 200x and 50x coverage, respectively. Genomic DNA was isolated using the QIAamp DNA Mini kit (Qiagen). Samples were sequenced (HiSeq 4000 instrument, Illumina Inc) in paired-end mode on Illumina (2×101 bp) using TrueSeq DNA exome kit (v.6) (Illumina Inc.). Paired-end reads were aligned to the human reference genome sequence GRCh38 using BWA–MEM (V0.715-r1140) (31). Deduplication and alignment metrics were performed using Picard tools (v2.12.1). Local realignment was performed around indels to improve SNP calling in these conflicting areas with the IndelRealigner tool. Recalibration to avoid biases was performed following the Genome Analysis Toolkit (GATK) Best Practices (32). Single mpileup files were generated from paired bam normal/tumor using samtools mpileup (v1.6). Mutation calling and computation of somatic p-values (SPV) was performed on mpileup output files using Varscan (v2.3.9)(33) to WES data from tumor and patient-matched normal samples with a minimum coverage of 10x. Quality control metrics were assessed using FastQC (v0.11.2)(34) before and after the alignment workflow and reviewed to identify potential low-quality data files.

### Single cell data processing

Sequencing data was processed with 10X Genomics Cell Ranger v2.1.1 with respect to GRCh38-1.2.0 genome reference to obtain UMI-corrected transcript raw gene expression count tables, BAM files and the BCR contig files.

To generate single cell allelic transcript counts we used a custom made script to identify reads intersecting WES-based mutated positions. For each read, to classify the allele we identified the single nucleotide overlapping the mutated base. To obtain transcript counts we used the unique molecular identifiers (UMIs) associated with the reads.

We used the vireo function from cardelino package v0.4.2 to construct clusters of cells sharing the same genotype. As input we provided allelic counts for the positions likely to differ between the subjects and not mutated between FL and stromal cells. For further processing we selected cells assigned to a single subject at the threshold of 0.75. Once the clusters of cells sharing the same genotype were identified, we assigned them to patients by comparing the cluster consensus genotype with the patient-labeled genotypes obtained from WES.

IMGT/HighV-Quest (35) was used for high-throughput BCR analysis and annotation of all_contig.fasta file (35). IMGT/HighV-Quest output data was filtered for productive and rearranged sequences and FL cells with identical BCR heavy chains were considered unique clones within the malignant cell population and were annotated with unique clonotype identifiers. R-package ‘vegan’ was used to calculate Pielou’s index of evenness for clonotype distribution.

### Phylogenetic analysis

For each subject, we first identified common mutations that can be found in both WES data and scRNA-seq data. Next, we used FALCON-X with default parameters for estimation of allele-specific copy numbers. As a verification, we compared the results of FALCON-X with those of GATK CNV analysis pipeline, and confirmed that the two approaches gave similar results. Finally, we run Canopy (9), providing the estimated major and minor copy number, as well as the allele-specific read counts in the tumor and matched normal WES data as input. Taking advantage of a Bayesian framework, Canopy estimates the clonal structure of the tumor for a pre-specified number of clones. Choosing between trees with the number of clones from 2 to 4, for both subjects, the BIC criterion used by Canopy suggested trees with 4 clones as the best solution. For further analysis, for each subject, we selected the top tree returned by Canopy.

### Mapping BCR clonotypes to tumor clones using CACTUS

Below we introduce a probabilistic model, CACTUS, for mapping a given set of cell clusters to tumor clones based on the mutation matching between the cells in clusters and the clone genotypes (Fig. 6). In this analysis, the clusters corresponded to the BCR clonotypes. For each subject, CACTUS was run for the top Canopy tree for a maximum of 20000 iterations, with 10 different starting points. For the sake of comparison, cardelino was applied with the same setup.

**Fig. 6.**
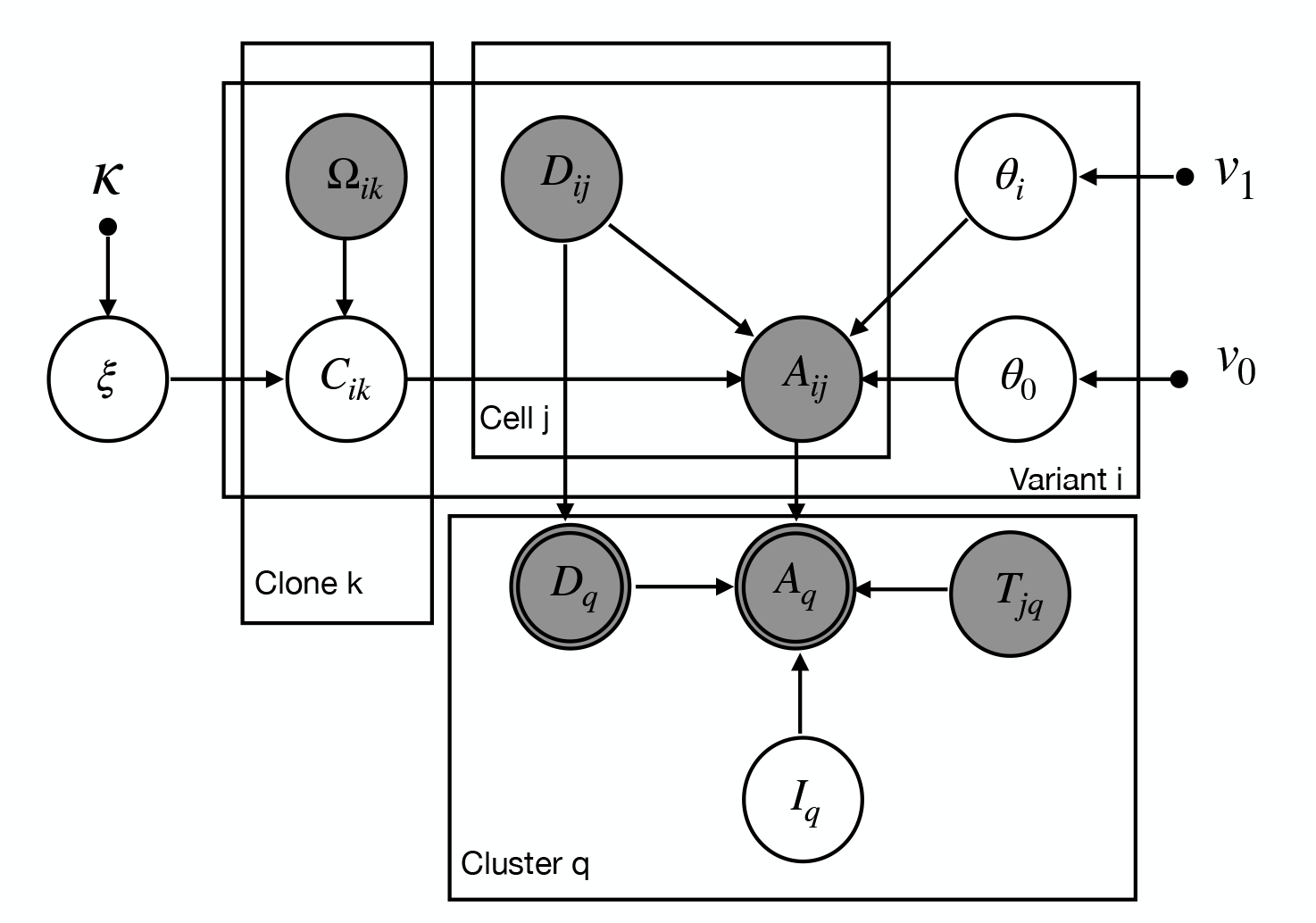
The graphical model representation of CACTUS. Circle nodes are labeled with random variables in the model. Arrows correspond to local conditional probability distributions of the child variables given the parent variables. Observed variables are shown as grayed nodes. Double-circled nodes are deterministically obtained from their parent variables. Small filled circles correspond to hyperparameters.

CACTUS is a direct extension of cardelino (21), accounting for cell clustering, with the assumption that cells in the same cluster belong to the same clone. Let *i* ∈ {1,…,*N*} index mutation positions, which can be identified both in bulk DNA sequencing and single cell RNA seq data (see above). We assume we are given at input a set of *K* tumor clones, indexed by *k* ∈ {1,…,*K*}. Each tumor clone is represented by its genotype and prevalence in the tumor population. The input clone genotypes are represented by a binary matrix Ω_*i,k*_ with entries equal 1 if the mutation *i* is present in clone *k* and 0 otherwise. We are also given an independent clustering of single cells, where each cluster *q* ∈ {1,…*Q*} contains a number of cells and the clusters are assumed not to overlap. Let *j* ∈ {1,…,*M*} index cells. The clustering is given by a binary matrix **T**, with *T_j,q_* = 1 if cell *j* is in cluster *q* and 0 otherwise.

We are interested in assignment of the given cell clusters to the given clones. The clone assignment of each cluster *q* is represented in the model by a hidden variable *I_q_* with values in 1*,…, K*. We assume a uniform prior for *I_q_* and set 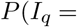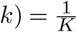. Alternatively, the prior could depend on the prevalences derived from the evolutionary model.

We assume that the input genotypes can contain errors with error rate *ξ*. The prior distribution for the error rate is parametrized by *κ* = (*κ*_0_*, κ*_1_) and is set to *P*(*ξ|κ*) = Beta(*ξ*; *κ*_0_*, κ*_1_). We define a hidden random variable *C_i,k_* denoting the true (corrected) genotype of clone *k* at variant position *i*, with

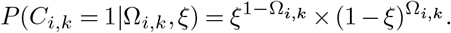

Let matrix **A** with elements *A_i,j_* denote the observed count of unique transcripts with alternative (mutated) nucleotide mapped to position *i* in cell *j*, and matrix **D** with elements *D_i,j_* denote the total unique transcripts count mapped to that position in that cell. Let *θ_i_* denote the success probability of observing a transcript with alternative nucleotide at a position *i* in a cell that carries this mutation, and *θ*_0_ the success probability of observing a transcript with alternative nucleotide in a position that is not present in the cell. The distribution of the observed read counts then becomes

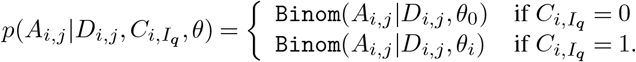

We assume Beta priors on the *θ* parameters

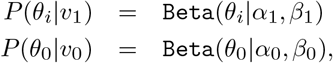

where *v*_1_ = (*α*_1_,*β*_1_) and *v*_0_ = (*α*_0_,*β*_0_). We denote *v* = (*v*_0_, *v*_1_).

Let *A_q_* be the matrix of alternative allele counts for cells contained in cluster *q*, at mutated positions, i.e., *A_q_* = (*A_i,j_*)_*j∈q,i*=1,…*N*_, and let *D_q_* = (*D_i,j_*)_*j∈q,i*=1,…*N*_. Since we assume the observed read counts at the different positions and different cells are independent, we have

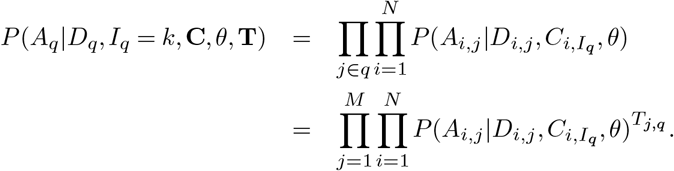

### CACTUS model inference

We use Gibbs sampler, a Markov chain Monte Carlo (MCMC) algorithm for generating samples from the posterior distribution. We iteratively sample each hidden variable which is conditionally independent given the rest of the hidden variables in the model. The hidden variables in CACTUS include the cluster assignment matrix **I**, the success probabilities of observing a transcript *θ* = (*θ*_0_*, θ*_1_*,…, θ_N_*), the corrected genotype matrix **C**, and its error rate *ξ*. We describe the sampling steps for the full joint distribution of these hidden variables in the following.

### Sampling clone assignment of clusters I_q_

We sample cluster-to-clone assignment variable *I_q_*, given the Markov blanket of *I_q_* in the graphical model (Fig. 6)

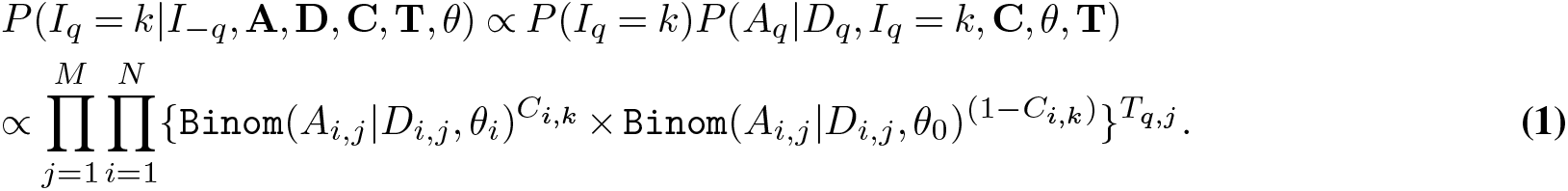

### Sampling success probabilities of observing a transcript θ

Similarly, we sample *θ_i_*, 0 < *i < N* from the posterior probability

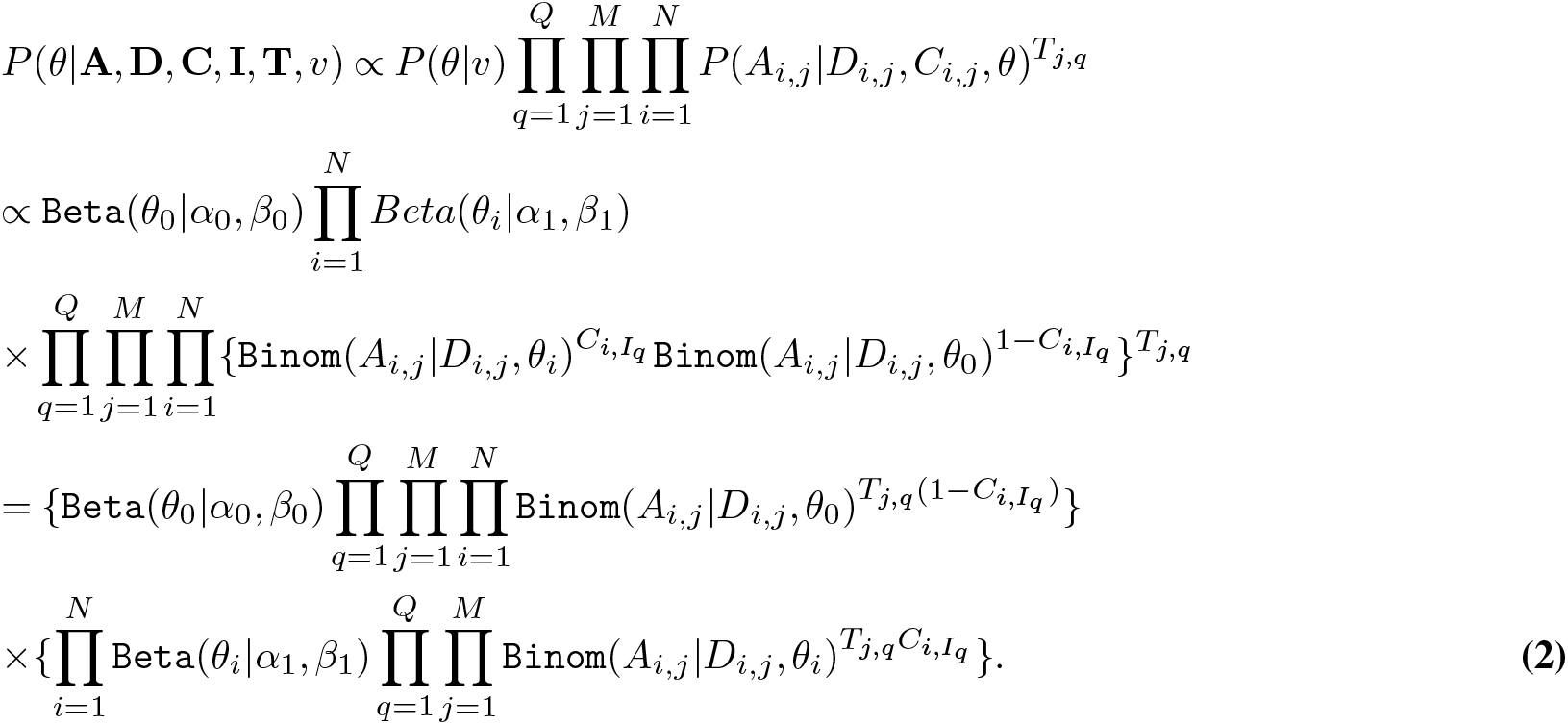

Using the Beta-Binomial conjugacy, *θ* is sampled from the Beta distribution

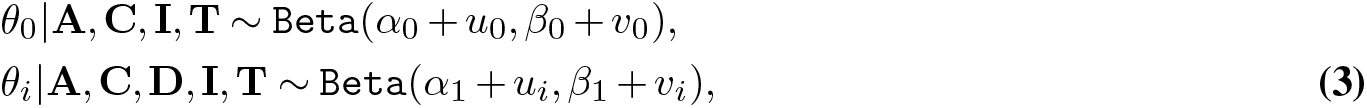

for *i* ∈ {1,…,*N*}, where

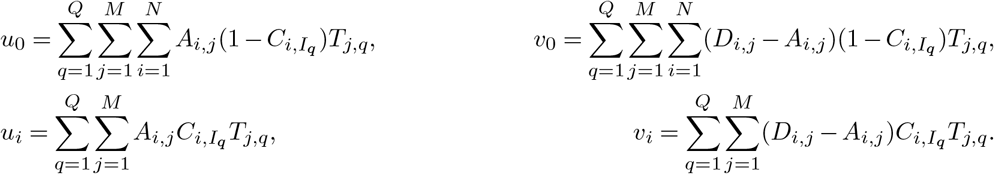

### Sampling the corrected genotype matrix C

The corrected genotype matrix **C** is sampled using

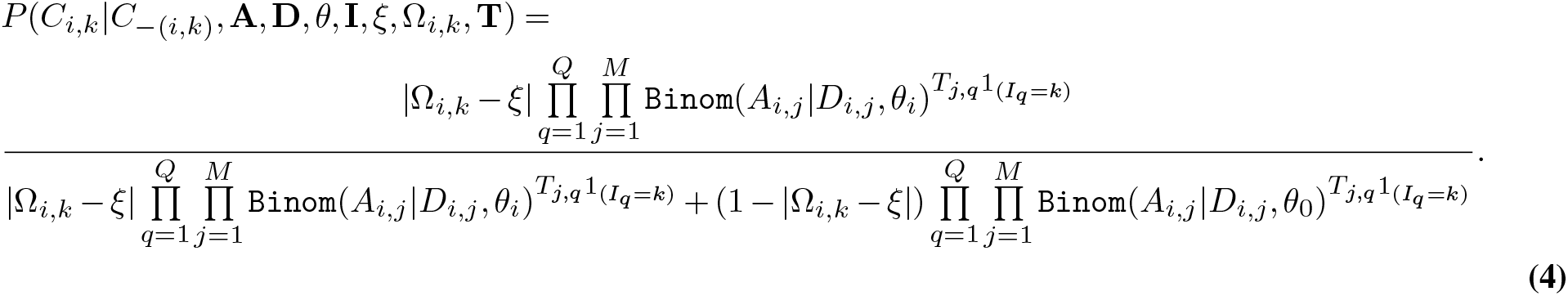

### Sampling the error rate ξ

We can compute the distribution of the error rate *ξ* having the corrected genotype matrix *C*, as well as the input genotype matrix Ω and hyperparameters *κ* as follows,

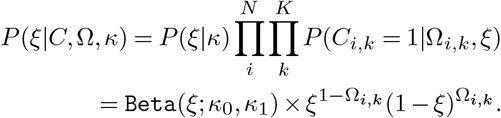

From the Beta-Bernoulli conjugacy we obtain

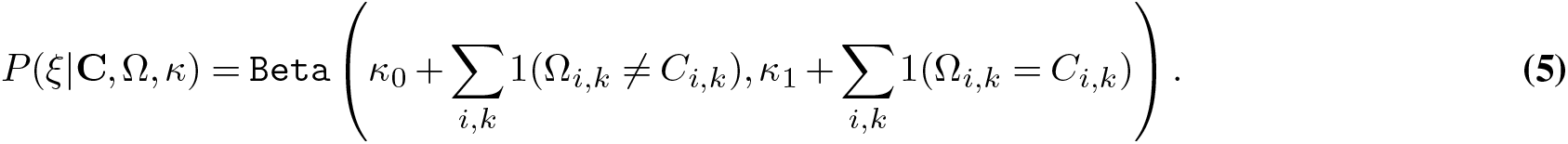

Finally, the Gibbs sampling algorithm for CACTUS was derived as a straightforward modification of the algorithm used for cardelino (21). In the algorithm, *I_q_* is iteratively sampled using Eq. (1) for *q* = 1, … *Q*, *θ_i_* for *i* = 1, …, *N* is sampled using Eq. (3), *C_i,k_* for *i* = 1, …, *N* and *k* = 1*, … K* is sampled using Eq. (4), and *ξ* is sampled using Eq. (5).

## Declarations

### Ethics approval and consent to participate, consent for publication

Lymph node biopsies were collected from patients after approval by the institutional review board (IRB) of the Leiden University Medical Center, Albinusdreef 2, 2333 ZA Leiden, The Netherlands, according to the Declaration of Helsinki. Prior written informed consent was obtained from all patients to investigate materials and to publish data and case details.

### Availability of data and code

The data used to produce the results presented in this publication, as well as the code implementing CACTUS are available at https://github.com/LUMC/CACTUS.

### Competing interests

The authors declare that they have no competing interests.

### Funding

This project has received funding from the European Union’s Horizon 2020 research and innovation programme under the Marie Skłodowska-Curie grant agreement No 766030. ES acknowledges the support from the Polish National Science Centre OPUS grant no 2019/33/B/NZ2/00956.

### Author’s contributions

S.D.S and E.S. developed the probabilistic model. S.D.S implemented the model, phylogenetic analysis, and carried out the application of the model and benchmarked an alternative method, supervised by E.S. S.D.S, D.C., and H.M. performed copy number calling in WES data. S.M.K. performed clustering of single cells to subjects and supervised primary data analyses. J.S. conducted mutation calling in WES data. R.Mo. performed single cell data sample deconvolution. R.Me. conducted alignment of scRNA reads. S.K. carried out exome and scRNA sequencing. H.V. provided patient samples and data. C.A.M.v.B. conceived and planned the experiments, carried out sample preparation and identification of BCR clonotypes. E.S., C.A.M.v.B., S.M.K. and S.D.S. conceived of the study and wrote the paper.

## Acknowledgements

The authors wish to thank Flow Core Facility operators Guido de Roo and Edwin de Haas for exquisite flow cytometric cell sorting and Emile Meijer of the Leiden Genome Technology Center for excellent preparation of single cell sequencing libraries.

## Supplementary Figures

**Fig. S1.**
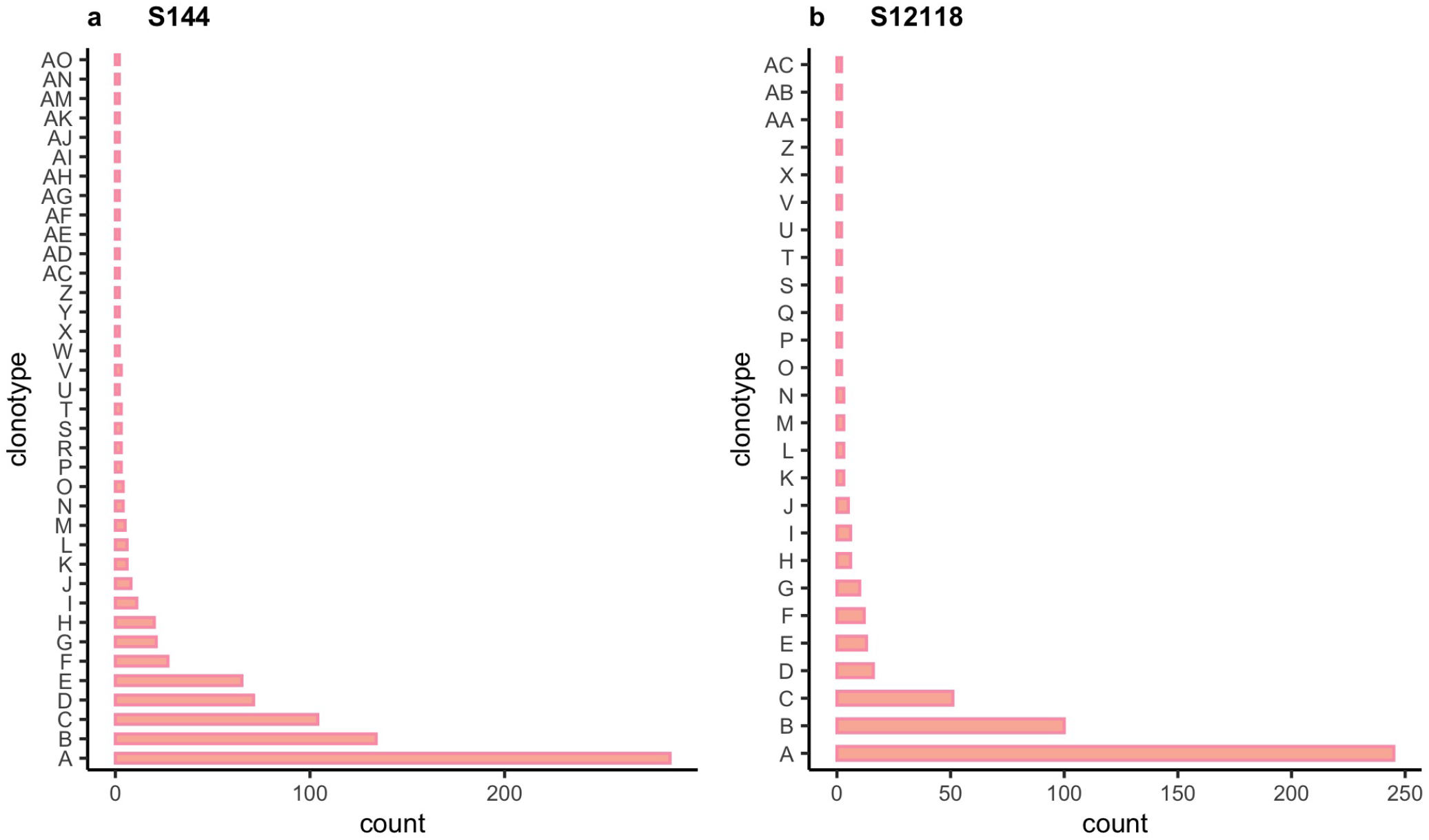
**Number of cells belonging to each BCR clonotype** for (a) subject S144 and (b) subject S12118. Only clonotypes with more than one cell are plotted. There are 442 clonotypes consisting of one cell for subject S144 and 299 clonotypes with one cell for subject S12118.

**Fig. S2.**
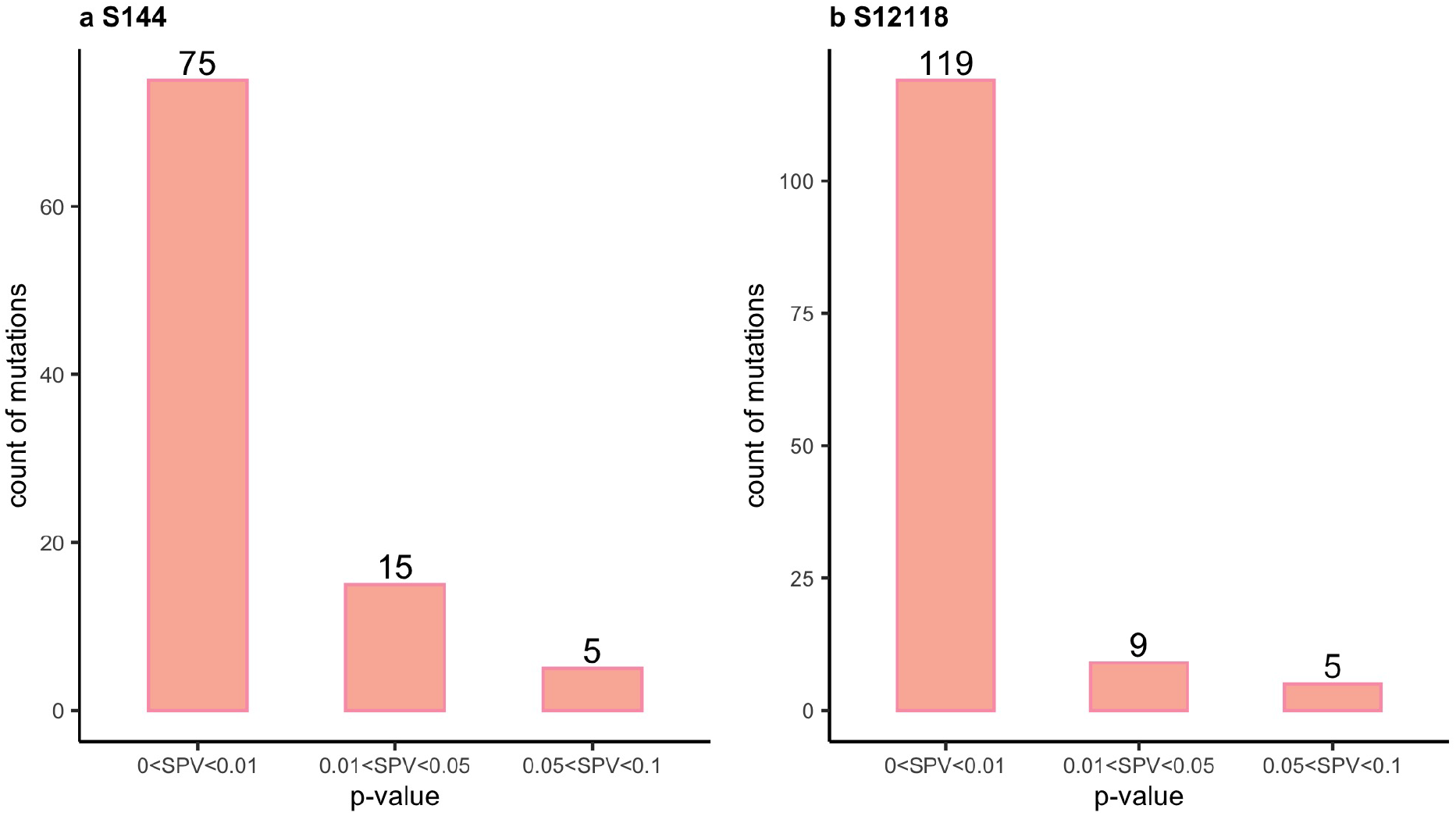
Somatic variant p-values for mutations that were shared between WES and scRNA-seq data. Somatic p-values (SPV) are grouped into three intervals: [0, 0.01], [0.01, 0.05], [0.05, 0.1] for (**a**) subject S144 and (**b**) subject S12118.

**Fig. S3.**
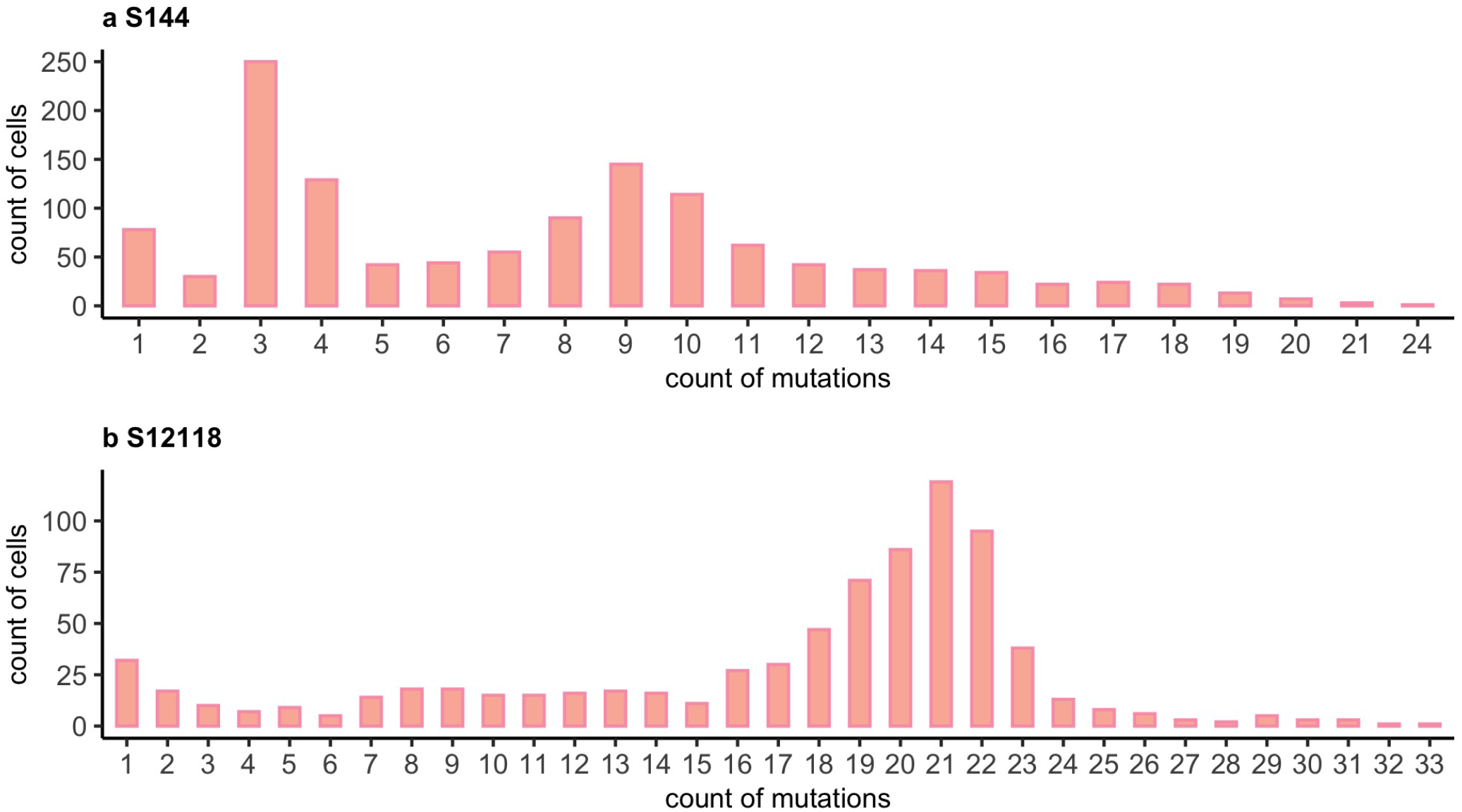
**Numbers of cells (y-axis) with specific mutation counts (x-axis)** for the mutations that were detected both in WES and scRNA-seq data for (**a**) subject S144 and (**b**) subject S12118.

## Notes

https://github.com/LUMC/CACTUS/

### Competing Interest Statement

The authors have declared no competing interest.

